# Movement path as an ethological lens into interval timing

**DOI:** 10.1101/2025.01.24.634414

**Authors:** Fuat Balcı, Varsovia Hernandez, Ahmet Hoşer, Alejandro León

**Author notes:** Correspondence concerning this article should be addressed to: Fuat Balci 50 Sifton Rd, Biological Sciences Building, Room 222, Winnipeg, R3T 2M5, Winnipeg, MB, Canada., Alejandro León Universidad Veracruzana, Centro de Investigaciones Biomédicas. Dr. Luis Castelazo Ayala s/n, C.P. 91190, Col. Industrial Á nimas Xalapa, Veracruz, México. Building “T”.

## Abstract

Interval timing behavior is traditionally investigated in operant chambers based on the very focal responses of the subjects (e.g., head entry to the magazine, lever press). These measures are blind to the movement trajectory of the animals and capture only a tiny segment and sometimes an idiosyncratic component of the animal’s behavior. In other words, the state of the temporal expectancy is not observable at every time point in the trial. On the other hand, in nature, temporal expectancies guide actions in a much more complex fashion. For instance, an animal might approach a food patch at different degrees as a function of the expected time of food availability (e.g., nectar collection behavior). The current study aimed to investigate interval timing in a more naturalistic fashion by analyzing the movement trajectory of rats in *fixed time* (*FT-30s*) vs. *variable time* (*VT-30s*) schedules in modified open field equipment. We observed a temporally patterned movement in *FT* but not in a *VT* schedule. In the *FT* schedule, rats moved away from the reward grid after consuming the presented water and were farthest from it for around 15 seconds, after which they started to approach the reward grid again. There was no such temporal patterning of movement trajectory in the *VT* schedule. Temporal control in the *FT* schedule was stronger in the *Wall* compared to the *Center* condition. Our results show that movement trajectory may reflect instantaneous temporal expectancy.

## 1 Introduction

Cognitive functions are adjusted to particular ecological contexts with critical spatial (e.g., location of food sources, risks, etc.) and temporal features (e.g., periodicity of the resources). For instance, spatial learning is adaptively embedded in the foraging behavior of animals. Imagine a foraging honey bee that searches, identifies, and learns a food source, which it will revisit and communicate to nest-mates via its waggle dance (Von Frisch, 1967). Highlighting the relevance of temporal features in foraging, hummingbirds revisit food sources at different delays depending on the time it takes them to replenish (Henderson et al., 2006). These clearly demonstrate how spatial and temporal features are relevant to naturalistic behaviors and the organisms’ adaptiveness. The study of cognitive function as part of ecological context constitutes the primary premise of the cognitive ethological approach (Vauclair, 1997).

In contrast to this naturalistic approach to animal cognition, the conventional psychological approach to studying animal behavior isolates the function of interest from its relevant ecological setting to maximize experimental control for a more direct quantification and investigation (see Timberlake, 2012). A few undertakings acknowledged this meta-theoretical gap and attempted to overcome it by integrating behavioral testing in the living environment of single-housed animals (Balzani et al., 2018; Gallistel et al., 2014) to eliminate artificial segmentation of test periods or testing group-housed animals (e.g. IntelliCage, Kiryk et al., 2020) that allowed studying even factors such as competition (Benner et al., 2015). Relatedly, Agostino and Golombek (2022) have recently raised the need for a more naturalistic interpretation of the timing function and designing more naturalistic settings to investigate the timing function in the laboratory. These are notable empirical and meta-theoretical developments for naturalizing behavioral testing in general and the study of interval timing specifically.

Another gap that limits the ecologically relevant characterization of behavior relates to studying behaviors over a single dimension of interest while minimizing the uncertainty of task parameters in terms of other ecologically relevant dimensions. The conventional study of interval timing behavior, which refers to organizing responses around seconds-to-minutes long intervals (Buhusi & Meck, 2005), constitutes a good example of this limitation. Specifically, the interval timing behavior of lab animals typically requires subjects to anticipate reward delivery at a designated hopper (typically placed on a wall) or serially ordered hoppers associated with increasingly shorter or longer delays (e.g., Gallistel et al., 2004; Roberts, 1981).

As argued above, this approach does not sufficiently capture the complexity of timing behavior in naturalistic settings as it isolates the temporal aspects of behavior from its spatial features. To our knowledge, only a handful of studies have (albeit reduced in complexity) considered spatiotemporal characteristics of reward anticipation. From these, Machado and Keen (2003) tested pigeons in the long box where one side of the box was associated with the short and the other side of the box was associated with the long interval (spatial feature was reduced to one dimension). They found a stereotypical response pattern where pigeons first approached the “short side” and then switched to the “long side” if the reward was not present at the “short side” after the short delay; the latency to leave the “short side” was under the control of short latency (the time at the short side and arrival at the short side were negatively correlated).

A similar pattern was observed in the mice tested in the timed switch task, where the operanda were placed on the same wall of the operant chamber (e.g., Balci et al., 2008, 2009). The temporal periodicity of behavior by movement was also demonstrated in a task where the stimulus-reward contingencies were reversed mid-session, and the animals exhibited anticipatory errors based on the timing of the contingency reversal (Soares et al., 2024). These studies demonstrated timing behavior but over a reduced spatial relation (reduced to a movement over one dimension) and thus fall short of capturing the spatiotemporal complexity of behavior.

The movement trajectory of the organisms (e.g., mice and rats) as a function of two-dimensional space (relevant for lateral locomotor movements of non-flying animals) and time can provide a more in-depth understanding of the states of expectancy and complexity of anticipatory actions. There are a few examples of this unified approach in human research. For instance, McKinstry et al. (2008) showed that the spatial extent and temporal dynamics of reaching movements reflected the state of high-level cognition (i.e., competition in decision-making). Similarly, Maldonado et al. (2019) showed that the dynamics of the computer mouse path shed light on the underlying decision processes.

The current study aimed to investigate how the movement trajectory of rats elucidates the spatiotemporal modulation of reward expectancy based on a secondary analysis of published data (Hernández et al., 2023; León et al., 2020). Rats were tested in a modified open-field system that contained multiple grids as potential locations for reward delivery. The water reward was delivered after either fixed (*Fixed Time* [*FT*] schedule 30”) or variable intervals (*Variable Time* [*VT*] schedule 30”) in different experimental groups at one of the grids. The *VT* schedule was the control condition for the temporally predictable schedules (*FT*). The location of the active grid was altered within subjects and between different phases (*center* and *wall* [peripheral] location — order counterbalanced). We predicted that rats would become closer to the reinforcement location as the time of reinforcement delivery closed in (e.g., *> FT/*2) compared to the *VT* schedule.

## 2 Method

### 2.1 Subjects

Subjects were thirteen experimentally naive Wistar rats (10 female) acquired from the Animal Facility of the Autonomous University of the State of Hidalgo, Mexico, experimentally naive. At the start of the experiment, the rats were 3.5 months old and individually housed. Rats were housed in a 12h-12h dark-light cycle and tested during the light cycle of each day. They were subjected to a 23-hour water restriction with a 30-minute access period at the end of each session while the food was readily available in their home cages. Daily sessions lasted 20 minutes, and they were conducted seven days a week. All procedures complied with university regulations for animal use and care and the Mexican norm NOM-062-ZOO-1999 for Technical Specifications for the Production, Use, and Care of Laboratory Animals.

### 2.2 Apparatus

A Modified Open Field System (MOFS) was employed to assess the animal’s behavior. The MOFS comprised a chamber measuring 100 cm *×* 100 cm and was constructed with black Plexiglas panels on all four walls and the floor. Water could be presented in 100 locations, utilizing 0.8 cm diameter holes spaced 0.95 cm apart. The water dispenser, developed by Walden Modular Equipment®, operated on a servo system and was placed either near the wall (Wall Condition, coordinates 95, 45) or in proximity to the center of the MOFS (Center Condition, coordinates 45, 45). The central location of the water dispenser was positioned slightly closer to the right wall than the left wall, as none of the 100 water delivery locations were situated precisely at the center of the floor. The water dispenser released 0.1 cc of water, delivered through a water cup that emerged 0.4 cm above the floor of the MOFS via one of the holes. The MOFS was illuminated by two low-intensity lights (three watts) mounted above the chamber and positioned on opposite sides of the room, providing 9 lux of illumination to the chamber area. Following water delivery, the water was accessible for three seconds for consumption. A textured 9 cm *×* 9 cm black patch was 3D-printed with 16 dots per sticking out and placed 5.5 cm from the water dispenser for identification. The MOFS was cleaned with 70% isopropyl alcohol between each experimental session. The rat’s location was tracked using an RGB camera and Walden Software, which tracked the rat’s position in *X* and *Y* coordinates at a temporal resolution of 5 Hz in real time.

### 2.3 Procedure

Data were obtained from two experiments originally reported by León et al. (2020) and Hernández et al. (2023). In each experiment, subjects were randomly assigned to one of two groups: 1) *Fixed Time* or 2) *Variable Time*. Subjects from all groups were exposed to two consecutive conditions in a different order (see Table 1). In each condition, water was delivered using an *FT* 30 s schedule or a *VT* 30 s schedule. Note that the *wall* reward condition more closely represents the study of timing behavior in conventional operant boxes. In contrast, the *center* reward significantly diverges from the standard configuration of operant chambers. The *VT* schedule comprised a list of 7 values (3 s, 7 s, 13 s, 21 s, 31 s, 47 s, and 88 s) based on Fleshler and Hoffman (1962) progression, from which one value was randomly selected without replacement. This list of values for the *VT* schedule was chosen such that their mean was equal to the *FT* schedule (i.e., 30 s) with substantial variability (a common rule was not used to present possible oscillatory patterns in the data). The water remained available for 3 s, after which the reinforcement schedule started. In the *Peripheral Condition*, the water dispenser was placed on the floor adjacent to one wall of the experimental chamber. In contrast, in the *Center Condition*, the dispenser was located at the center of the experimental arena. Each condition was tested over 20 sessions in 20 minute long sessions. Each group comprised three subjects, except for group *VT* 30^*′′*^ *Periphery-Center*, which had four subjects.

**Table 1:**
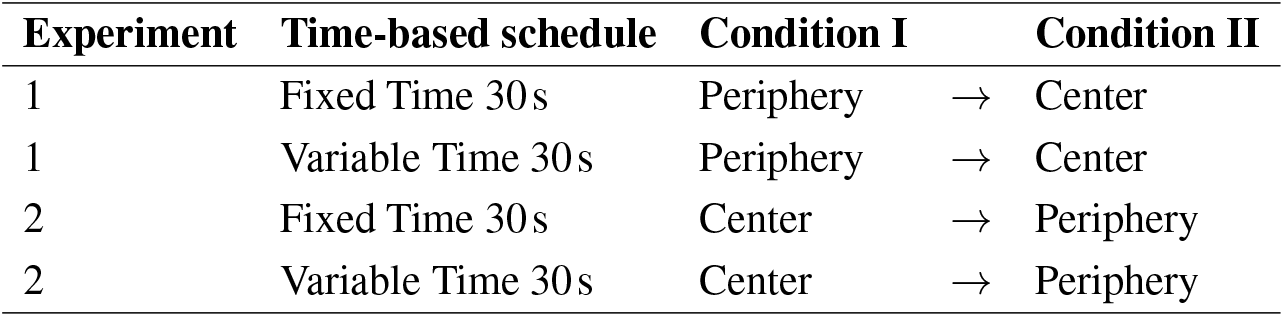
Experimental design.

| Experiment | Time-based schedule | Condition I | → | Condition II |
| --- | --- | --- | --- | --- |
| 1 | Fixed Time 30s | Periphery | → | Center |
| 1 | Variable Time 30s | Periphery | → | Center |
| 2 | Fixed Time 30s | Center | → | Periphery |
| 2 | Variable Time 30s | Center | → | Periphery |

### 2.4 Data Analysis

We conducted a secondary analysis of published data (Hernández et al., 2023; León et al., 2020). Data from the two experiments were pooled since the conditions were the same (except for the within-subject order of the reinforcement grid conditions). We quantified the Euclidean *distance* between the rats’ location in the open field and the reinforcement grid (central or peripheral) as a function of *time*:

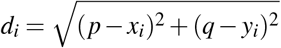

where (*p, q*) represents the coordinates of the reward (constant within a condition), (*x*_*i*_, *y*_*i*_) corresponds to the coordinates of the rat, and *di* is the distance between the rat’s location and reward location at frame *i*. The frame number was reset with reward delivery. For the *VT* schedule, all the data were used; the data collected in the 3 s schedule were included in the calculation of the means, while the data after that point onward were treated as “censored” and thus ignored as missing. This way of treating the data prevented any potential biases that would be introduced by differential censoring points in the *VT* schedule. These values were compared to the data collected from the full 30 s of the *FT* schedule.

We used the reciprocal of the Euclidean *distance* per frame (i.e., 1*/di*) as the dependent variable, which we predicted to increase as a function of *time* (following 15 s). On the other hand, there would not be an increase in *proximity* during the same period on the *VT* schedules (a flat or decreasing *proximity* is expected). We also tested whether temporal control over locomotion, particularly in *FT* schedules, was modulated by the location of reward delivery (middle of the open field or its wall). We analyzed the *proximity* (normalized per individual rat) data following 15 s using the following linear mixed effects model:

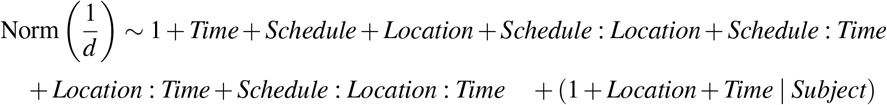

This statistical model included the three-way interaction between *Time >* 15 s, *Schedule* (*FT* vs. *VT*), and *Reward Location* (*Wall* vs. *Center*), including all lower-order interactions. Subjects were set as random variables for the intercept, slope, and reward locations (the best-fit model based on BIC scores compared to simpler models). The conditional *R*^2^ value of the model fit was 0.79 (marginal *R*^2^ = 0.11).

### Change Point Analysis

To estimate the *change points* in the location of rats, we applied the built-in findchangepts function of Matlab on the individual rat’s averaged *proximity* data. We used the linear variant of this function to find the points where the mean and the slope changed most abruptly. We allowed at most two *change points* in a series to capture the most prominent *change points*.

## 3 Results

### 3.1 Response Curve Analysis

Figure 1 shows the average normalized *proximity* values. The left column shows the data gathered from the *FT* condition, and the right column shows the data gathered from the *VT* condition. The visual inspection of this figure shows that after localization near the reward grid early in the trial and then disengagement from the consummatory behaviors, the *proximity* to the reward location increased linearly as the *time* of reinforcement was approaching in *FT* schedules. This temporal modulation of locomotion was more robust when the reward was delivered close to the *wall* than when it was delivered in the *center* of the testing equipment. On the contrary, rats became farther away from the reinforcement delivery location with *time* in the *VT* schedule. The initial peak (global maximum) after reward delivery at *time* zero corresponds to the localization of the rat around the reinforcement grid, and the downward righthand leg after the peak corresponds to the rat post-consumption moving away from the reinforcement grid.

**Figure 1.**
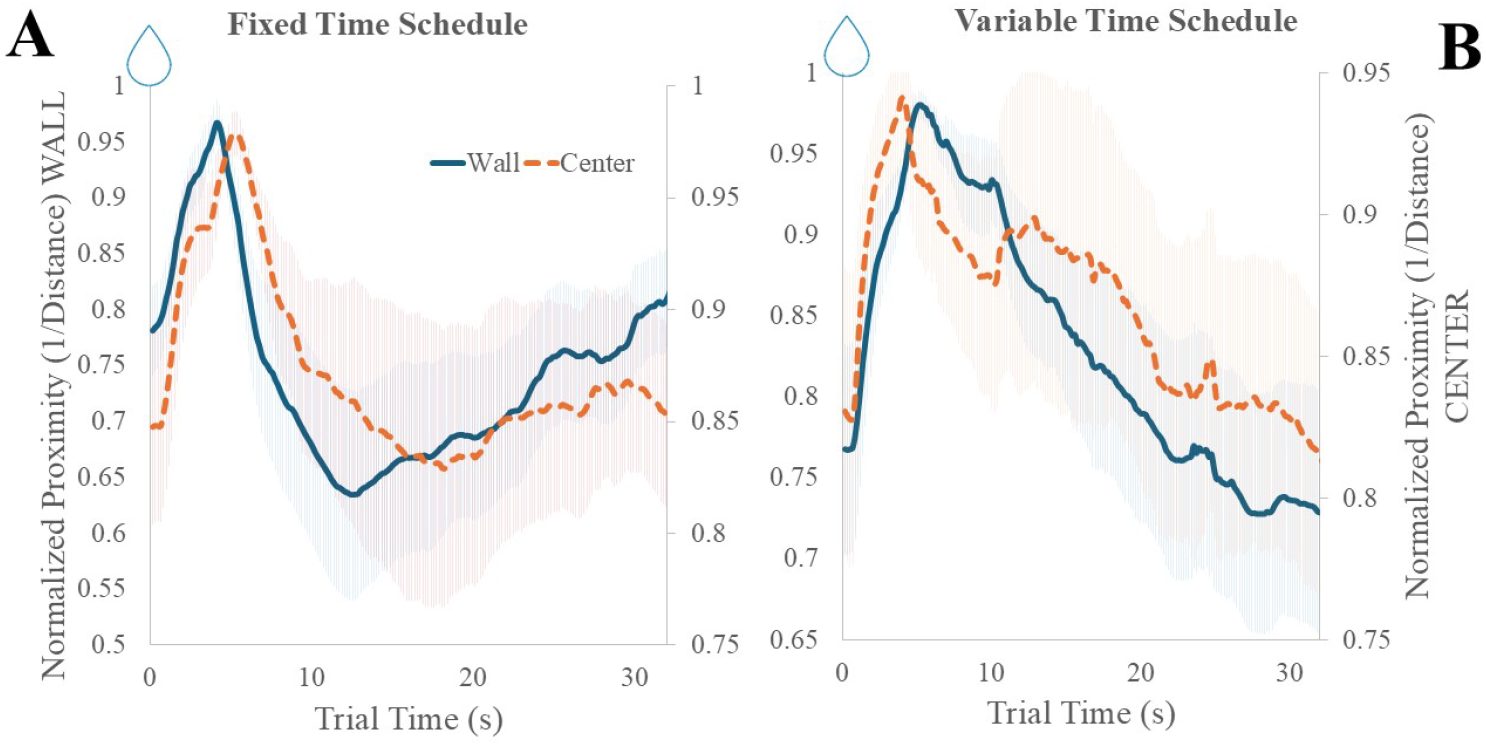
The average *proximity* (1*/Distance*) of rats to the reinforcement grid is separated for the *wall* (blue solid) and *center* (red dotted) reward locations. The left panel shows the data gathered from the *FT* condition, and the right panel shows the data gathered from the *VT* condition. The reward is delivered at time point zero (denoted by water drops). Left Panel: *FT* schedule. Right Panel: *VT* schedule. Note that the left y-axis shows values for the *wall*, and the right y-axis shows values for the *center* reward location. The *VT* schedule data are shown until 32 s for easier comparison to the *FT* schedule. When the average *proximity* to the reinforcement grid for the *VT* schedule was visualized up to longer intervals, there was still no temporal control. Error regions refer to the standard deviation of the mean.

The statistical analyses of the *proximity* data from 15 s (including the reward presentation period) onward corroborated these observations. There was a significant interaction effect of *time* with *schedule* (*F*(1, 2355.52) = 327.18, *p <* .001, Estimate *VT-FT* = −0.012); *proximity* to reward *location* increased with elapsing *time* in the *FT* and decreased in the *VT* schedule irrespective of the reward *location* condition. There was also a significant interaction effect of *time* with reward *location* (*F*(1, 2591.3) = 9.78, *p <* .01, Estimate *Center-Wall* = −0.002). Most importantly, there was a significant three-way interaction between *time, schedule*, and reward *location* (*F*(1, 2591.31) = 59.75, *p <* .001).

Figure 2 shows that the temporal control established by the *FT* schedule was much more prominent for the *Wall* than the *Center* condition. Simple effects analysis showed a significant positive slope for *FT* (*t*(12.3) = 9.09, *p <* .001) and a significant negative slope for *VT* (*t*(10.9) = −6.92, *p <* .001) in the *Wall* condition. The same results held in the *Center* condition; *FT*: (*t*(12.6) = 2.57, *p <* .05) and *VT*: (*t*(10.9) = −4.05, *p <* .01) in the *Center* condition.

**Figure 2.**
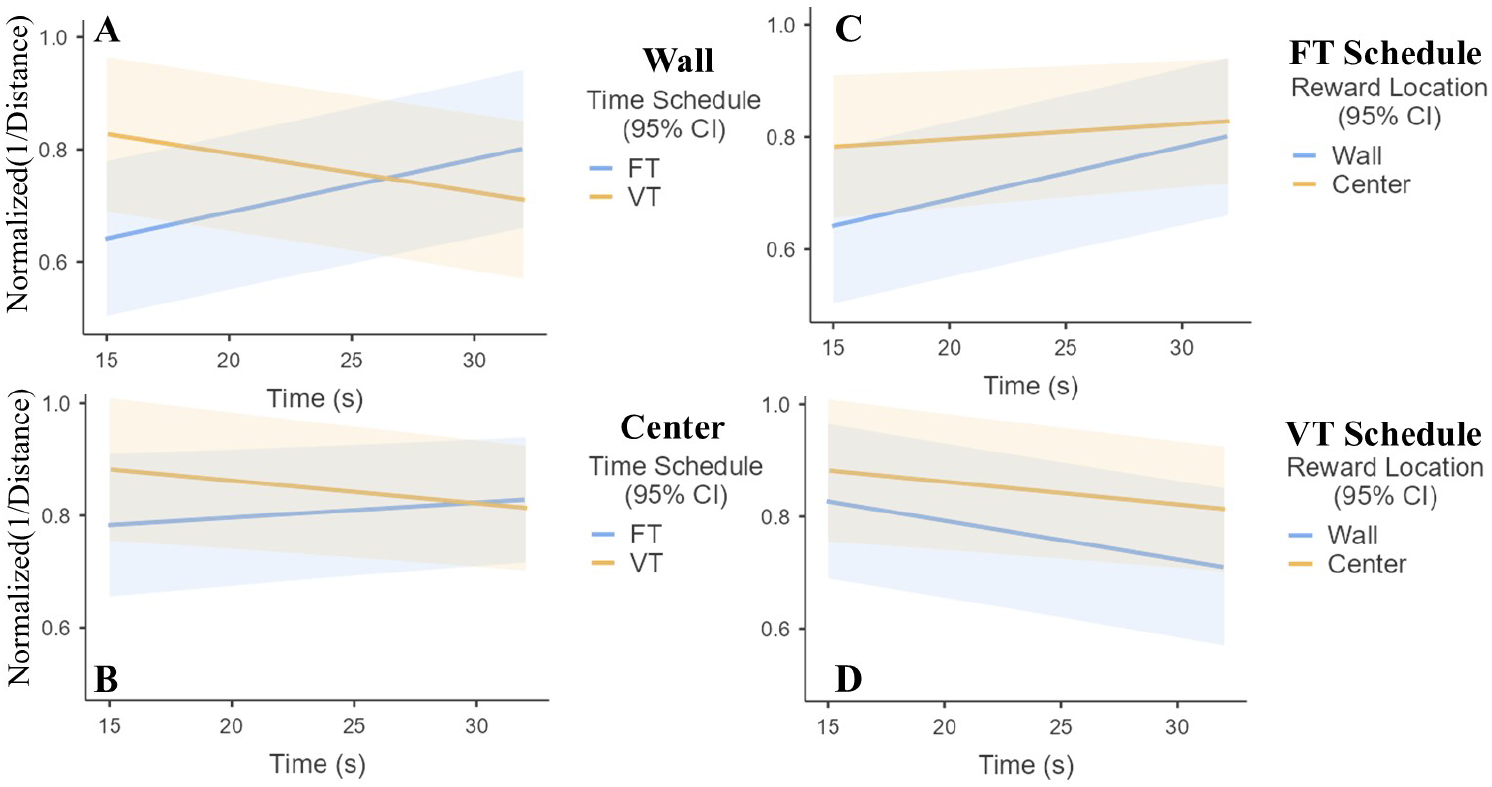
The modulation of normalized *proximity* is shown separately for the *wall* condition (top left panel) and the *center* condition (bottom left panel), as well as *FT* (dark line) and *VT* (light line). The same relations are separated by reinforcement schedule on the right panels (top right panel: *FT* schedule, bottom right panel: *VT* schedule). Shaded areas indicate the 95% confidence interval.

### 3.2 Change Point Analysis

To better characterize the patterns of changes in the *location* of the rats, we conducted a change point analysis of individual rats’ *proximity* data. If the temporal control over locomotion resembled the two-state pattern observed in traditional interval timing tasks in the operant box, one would expect an inflection point after which rats would approach the reinforcement grid. The visual inspection of Figure 1 indeed pointed to such a pattern where the global minima were observed between 12 s and 18 s in the *FT* schedule (akin to a bullish trend followed by cup formation in the technical analysis of financial markets), while the global minima were not reached within 32 s in the *VT* schedule.

The expected *proximity* pattern is a negative change point (disengagement from the reinforcement grid) followed by a positive change point (initiation of approaching behavior). Visual inspection of Figure 3 shows that this was indeed the case in 83% of the rats in the *FT* schedule while constituting 36% of the rats in the *VT* schedule. The visual inspection of Figure 3 also shows a higher degree of variability in the change points of rats in the *VT* schedule than the *FT* schedule, presumably because there was a common predictor of reward delivery in the latter condition. SOM shows the output of the change point analysis for individual rats.

**Figure 3.**
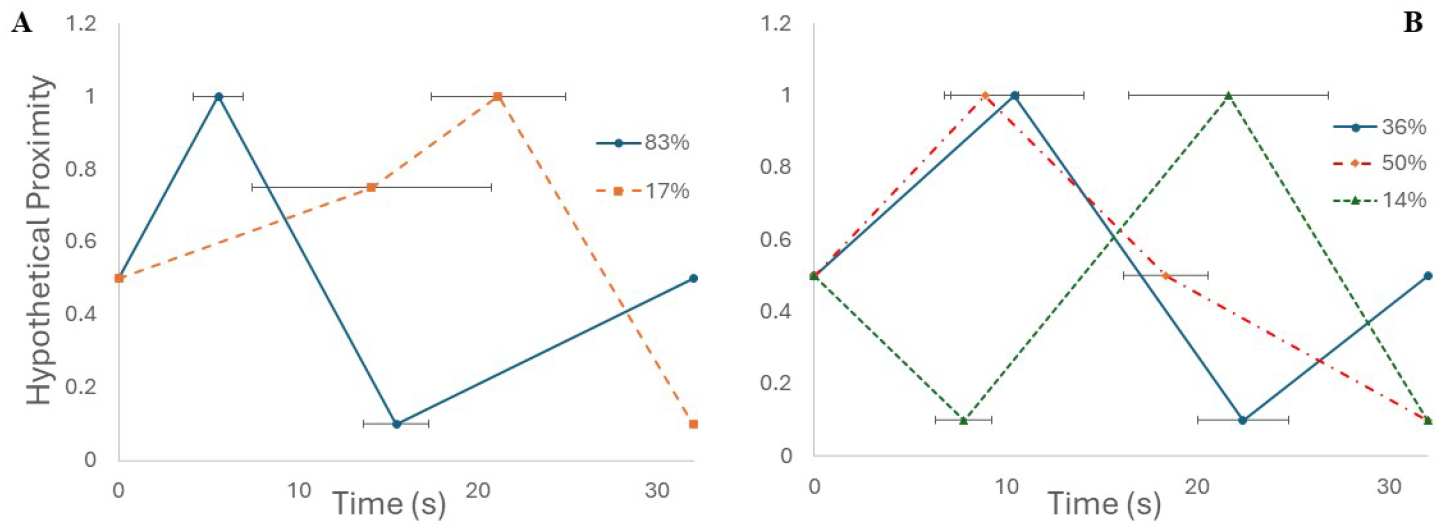
Different patterns of *proximity* dynamics in *FT* (A) and *VT* schedules (B). Two unique *proximity* patterns were detected in the *FT* and three unique *proximity* patterns in the *VT* schedule. Error bars are the SEM of change points.

## 4 Discussion

The current study demonstrates that the pattern of spatial location depends on the degree of predictability contained in time schedules according to which reinforcements become available in the environment. Specifically, we observed that when the reward was presented periodically every 30 seconds (*FT* schedule), rats changed their movement patterns with time to get closer to the reinforcement grid as the elapsed time approached the reinforcement delivery time. The continuous nature of locomotion affords a higher level of correspondence between temporal expectancy and spatial location. Notably, this level of analysis of spatiotemporal adjustment cannot be achieved using conventional operant tasks.

This pattern was more prominent when the reinforcement was presented peripherally (on the wall) than centrally. This difference might be because despite multiple sessions-long exposures to the test environment, rats avoided the central zone of the open field and preferred the peripheral zones, possibly as a form of preventing risk or anti-predation-like behaviors (Alstott & Timberlake, 2009; Martinez & Morato, 2004). Notably, the temporal modulation of spatial location was not present in the case of the *VT* schedule in which reward delivery was not predictable. The temporal modulation of spatial location in *FT* schedules can be due to the switch between the rats’ local and general search behaviors (former close to the water delivery, latter when the water delivery is distant in *time*). This is consistent with the transition between different modes outlined in Timberlake’s Behavioral Systems Theory (Freestone & Balci, 2019; Sanabria et al., 2019; Timberlake, 2012). Staddon and Simmelhag (1971) have also alluded to similar transitions between different response modes to explain the stereotypical responses under *FT* schedules.

In line with the abovementioned approach, the average response curves suggest an inflection point in the rat’s *proximity* to the reinforcement grid between 13-16 seconds. This observation is very similar to the transition from a low response rate to a high rate of responding that typically occurs around half of the delay to the reinforcement in *FT* and fixed interval-FI schedules (e.g., Gibbon et al., 1984). These breakpoints are referred to as start time, and they might be preceded by the approach behavior in the operant box, as shown in this paper. Our *change point analysis* of average response curves confirmed this observable pattern by detecting a positive *change point* toward the reinforcement grid around half of the *FT* schedule (with relatively low variability between rats). Note that the inflection points tended to be earlier (12 seconds) in the *Wall* condition (representing the setup in the conventional operant box) than in the *Center* condition. The earlier detection and lower variability of *change points* in the *Wall* condition could be due to the steeper modulation of *proximity* measures in the *Wall* condition. The initial prominent peak (i.e., global maximum), observed irrespective of schedules and conditions, denotes the approach behavior following the water delivery and thus cannot be attributed to temporal expectancy.

The present study expands the investigation of anticipatory behavior or interval timing beyond the conventional paradigms that are typically based on local discrete responses (e.g., nose poking, lever pressing). Our study provides robust evidence that animals adjust their continuous movements, such as displacement, to exploit the temporal statistics of the environment. This is significant because it allows for an ethologically-informed, more comprehensive timing framework and integrative approach to this fundamental function. It is important to note that animal movement, including approach-withdrawal patterns, is a fundamental response system with ecological relevance (Maier & Schneirla, 1964) and is a ubiquitous feature shared across diverse taxa. Therefore, studying temporal adjustments based on the movement trajectory of organisms can provide a plausible framework for a comparative approach to interval timing beyond specific response capabilities (e.g., manipulation) and response topographies in different taxa. The displacement with respect to a region of interest, in particular, might mirror the animals’ internal state regarding elapsed *time*.

A key theoretical question that arises from our work is whether the spatiotemporal modulation was indeed an expression/externalization of temporal expectancy or whether these spatial non-focal responses may be behaviors that facilitate interval timing as adjunctive states (e.g., Killeen and Fetterman, 1988; Killeen and Pellón, 2013). These two alternatives constitute distinct hypotheses that cannot be readily compared based on the current dataset. Future work should consider experimental designs and setups that can distinguish between these alternatives (e.g., introducing unpredictable barriers or changing the *location* of the flagged reinforcement grid).

The modulation of temporal adjustments by the dispenser *location* deserves special mention. Albeit speculative in nature, one plausible explanation for the different temporal adjustments with the same schedule but different dispenser *locations* could be related to the inherent tendency of rats to avoid open zones as a potential risk (Alstott & Timberlake, 2009; Martinez & Morato, 2004). It is possible that risk zones alter temporal adjustments or that timing behavior is not manifested under risky situations. This possibility can be tested in future research by increasing the perception of predation risk. For instance, Whitaker and Gilpin (2015) showed that rats exposed to predator odor spend less time in the central zone of the open field. An alternative explanation could be based on geometrical features inherent in the experimental setup, such as fewer adjacent open zones to the dispenser in the peripheral than central reward delivery condition. Relatedly, the maximal *distance* that can be recorded in the *Center* condition (∼ 64 cm) is lower than in the *Wall* condition (∼ 105 cm). Finally, it could be easier to discern the reward *location* relative to some physical landmark, like the wall, which would enhance procedural/temporal control over behavior and reduce noise. Future work is needed to elucidate these issues.

One of the limitations of the current study is that the specific behaviors of the rats were not identified and quantified (e.g., grooming, sniffing, rearing). Identifying responses at such a granular level would be more informative regarding the change in behavioral states/modes as a function of trial time (e.g., Bayesian Behavioral Systems Theory - Freestone and Balci [2019] or Markov chains - Sanabria et al. [2019]). Future studies should provide a more granular qualitative classification of response forms (Staddon & Simmelhag, 1971). A plausible solution to overcome this limitation and further expand the investigation of temporal adjustments is using advanced behavioral tracking techniques, such as *Pose Estimation tracking* or motion capture (Datta et al., 2019; Mathis & Mathis, 2020). These tools can identify and classify behaviors with high precision and temporal resolution, allowing for a more detailed and comprehensive analysis of the temporal adjustments of animal behavior.

Additionally, future studies could explore the potential interactions between temporal and spatial features in guiding animal behavior. For example, future studies can introduce both temporal and/or spatial uncertainty to elucidate better the information processing dynamics that underlie the movement trajectory. Overall, the present study provides valuable insights into the modulation of animal movement by different levels of temporal regularity and highlights the importance of considering continuous behavior and ecological relevance in investigating interval timing behavior. The fact that similar patterns of behavioral expressions of temporal information are observed in our tasks as those in earlier tasks points to the strong nature of temporal control over behaviors (e.g., inflection points in response rate and proximity). Future studies could also include peak interval trials (e.g., omissions of reinforcement or extinction trials), which would facilitate the analysis of movement following the reward availability time and graphing them per response functions of movement to compare these data more directly to conventional interval timing data. The study’s limitations suggest potential avenues for future research to advance further our understanding of temporal adjustments of animal movement and their underlying spatiotemporal dynamics.

## 5 Declarations

### 5.4 Author Contributions

A.L. and V.H. collected the data; F.B., A.L., V.H., and A.H. conceptualized the data analysis; F.B. and A.L. analyzed the data; F.B., A.L., V.H., and A.H. wrote the manuscript.

### 5.5 Availability of Data and Material

Data and code are available for the reviewers. Upon publication, data will be available from the corresponding authors on reasonable request.

### 5.6 Ethics Approval

All data are approved by the Mexican norm NOM-062-ZOO-1999 for Technical Specification for the Production, Use, and Care of Laboratory Animals.

## 6 Supplementary Material

The Supplementary Material (SOM) includes additional figures supporting the results presented in the main text:

- **SOM Figure 1:** Change points (maximum two) determined based on individual rats’ normalized proximity data for *FT*.
- **SOM Figure 2:** Change points (maximum two) determined based on individual rats’ normalized proximity data for *VT*.

The Supplementary Material is available on the preprint platform.

