## Supplemental Online Material for "Movement path as an ethological lens into interval timing"

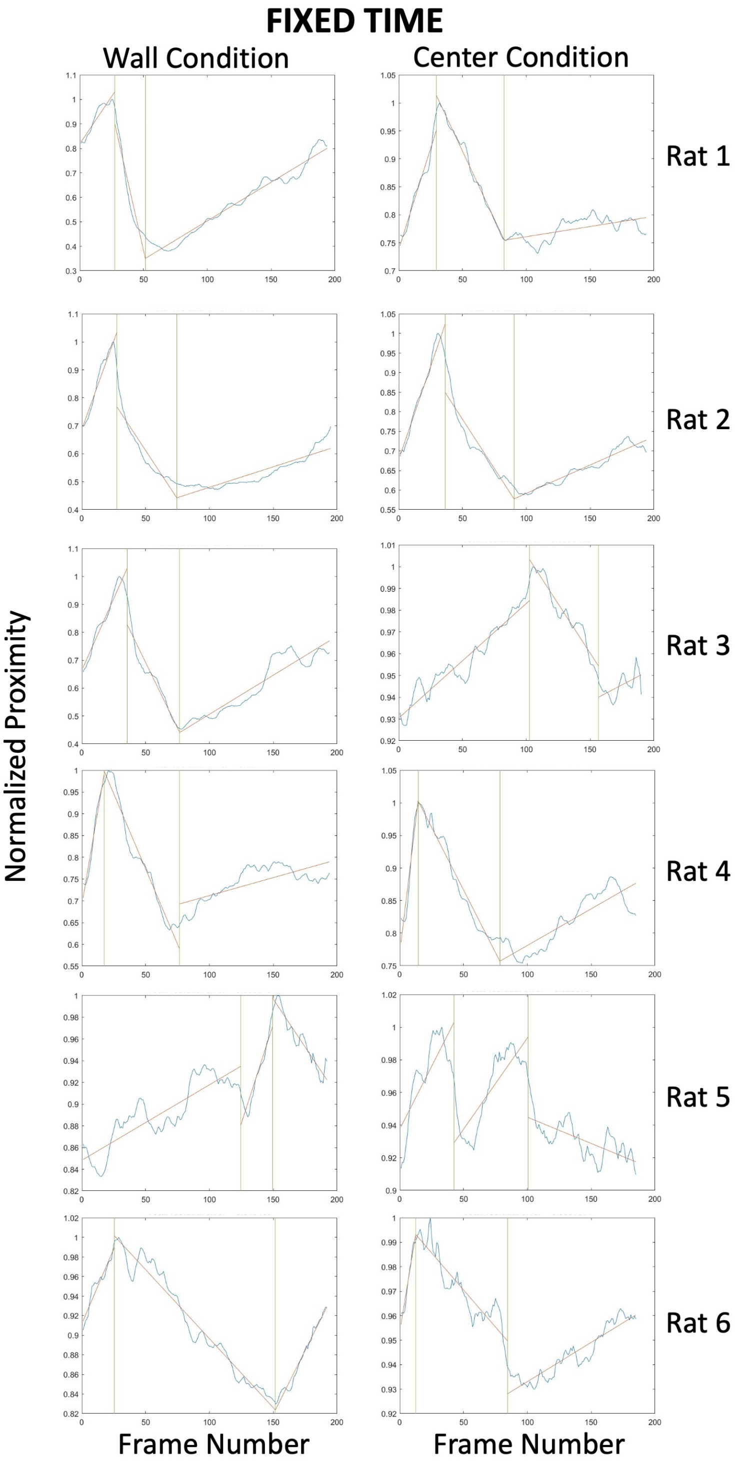


**SOM Figure 1.** Change points (maximum two) determined based on individual rats' normalized proximity data. The visual inspection of the plots shows a relatively consistent pattern for the FT (see Figure 3 of the main text).

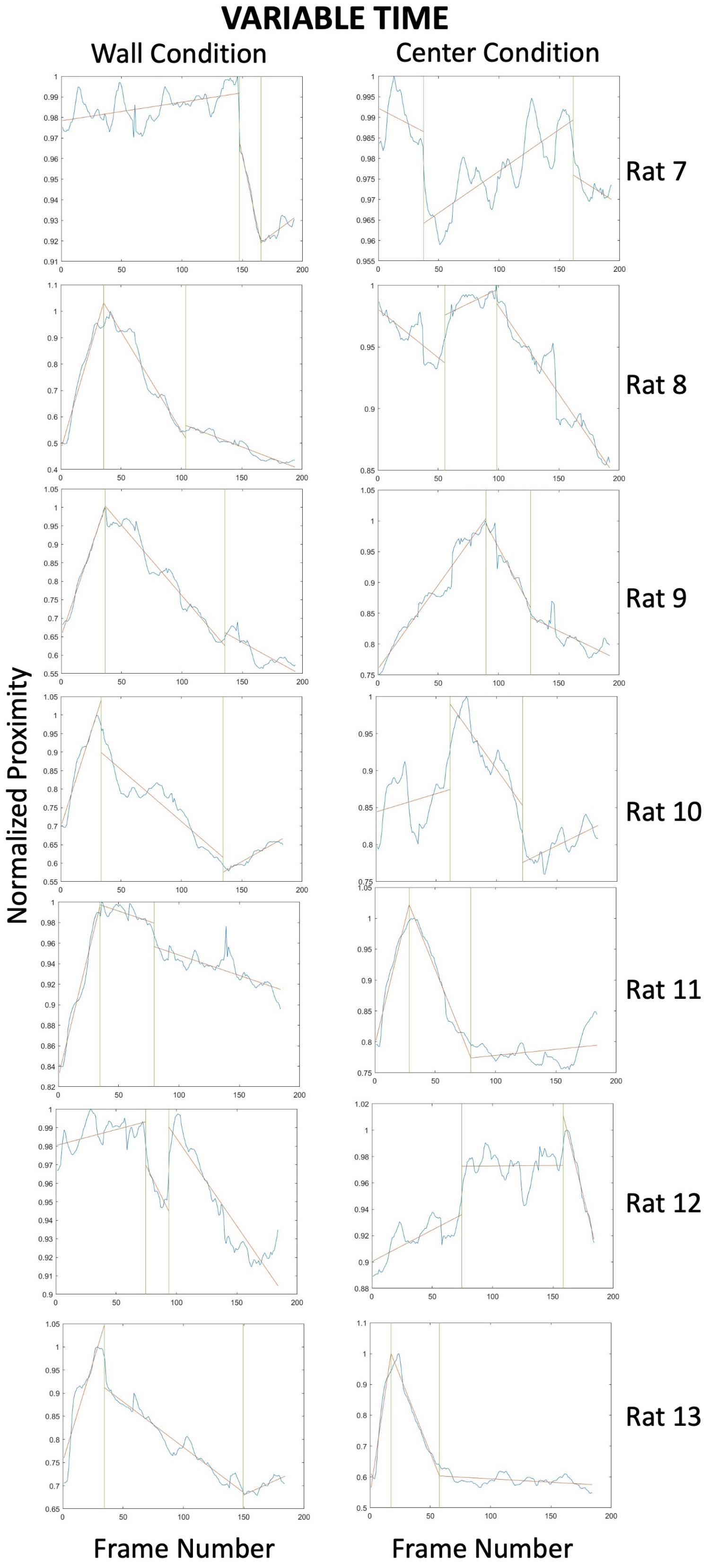


**SOM Figure 2.** Change points (maximum two) determined based on individual rats' normalized proximity data. The visual inspection of the plots shows more variable patterns for the VT schedule (see Figure 3 of the main text).
